# Developmental stage and level of submersion in water impact the viability of lone star and winter tick eggs during flooding

**DOI:** 10.1101/2024.05.18.594128

**Authors:** Maher Ramadan I. Alhawsawi, David A. Lewis, Ronja A. Frigard, Ellen M. Smith, Jaishna Sivakumar, Ajay M. Sharma, Adalynn R. Nantz, Chloe Sabile, Jasmine Kennedy, Rashi Loni, Gabrielle LeFevre, Akshita Vaka, Quinn Leanza, Melissa Kelley, Crystal L. Stacey, Richa A. Santhosh, Nathan Catlett, Tabitha L. Cady, Raaidh S. Rizvi, Zach Wagner, Pia U. Olafson, Joshua B. Benoit

**Author notes:** Author for correspondence: Joshua B. Benoit, Department of Biological Sciences, University of Cincinnati, OH 45221, USA., E-mail address (J.B. Benoit). authors contributed equally (Maher Ramadan I. Alhawsawi, David A. Lewis).

## Abstract

Female ticks deposit large egg clusters that range in size from hundreds to thousands eggs each. These clusters are immobile and restricted to a deposition site, usually under leaf litter and other debris. These sites can be exposed to periodic flooding, where the cluster of tick eggs can float to the surface or remain underneath organic debris entirely submerged underwater. Here, we examined the viability of egg clusters from winter ticks, *Dermacentor albipictus*, and lone star ticks, *Amblyomma americanum*, when partially or fully submerged in water and in relation to the developmental stages of the eggs. In general, egg clusters that were older and partially submerged had a higher viability than fully submerged, younger eggs. *A. americanum* was more resistant to water exposure between the two species. These studies highlight that egg clusters for certain tick species can remain viable when exposed to water for at least two weeks. These studies also suggest that distribution by flooding of egg clusters could occur for some species and flooding will differentially impact tick egg survival based on the specific developmental stage of exposure and species.

## Introduction

Ticks must survive between periods of feeding, which can consist of dry bouts as well as periods of water submersion. Mobile stages (i.e., larvae, nymphs, and adults) of multiple species have been evaluated for survival underwater and show extended periods of survival as these stages can absorb sufficient oxygen directly from the water due to their low metabolic rates (Lighton and Fielden, 1995; Fielden et al., 2011; Rosendale et al., 2019). The ability of tick stages to survive flooding is likely critical to many species as animal-tick interactions commonly occur in areas near water sources that may experience periodic flooding (Honzáková, 1971; Koch, 1986; Adejinmi, 2011; Giannelli et al., 2012; Bidder et al., 2019). Of importance is that the mobile stages can move to avoid flooding or climb upward, allowing behavioral avoidance or reduction in water exposure during flooding. Eggs, along with recently fed stages, are restricted to the site of deposition by the females or after feeding to where the feed stages settle to begin development to the next instar (Sonenshine and Roe, 2013).

Studies on the exposure of tick eggs to water sources is limited, but there is evidence of variability between and within species (Koch, 1986; Giannelli et al., 2012). In general, eggs remained viable when under water for less than five days, but viability was significantly reduced after 5-10 days (Koch, 1986; Adejinmi, 2011; Giannelli et al., 2012). Importantly, these studies focused on eggs that were counted and removed (either individually or as smaller clusters subdivided by a specific mass) from naturally laid egg clusters rather than examining how water submersion impacts egg viability as a group. This is critical as disturbing egg clusters reduces viability (Benoit et al., 2007; Yoder et al., 2012; Ramos et al., 2013). Furthermore, the egg stage, whether early or late in development, is important to stress tolerance (Ajayi et al., 2024), which was not previously examined in relation to water submersion.

In this study, we examined the viability of egg clusters in response to water submersion in two tick species, winter ticks, *Dermacentor albipictus*, and lone star ticks, *Amblyomma americanum*. As tick eggs, specifically when in clusters, float on the surface of the water, we measured viability of partially submerged (floating on the surface) egg clusters compared to those that are fully submerged in water. Furthermore, we have recently established that stress tolerance in tick eggs is dependent on the process of embryo and larval development within the eggs (Ajayi et al., 2024). As such, we determined whether or not the developmental stage of the tick eggs has a direct impact on egg survival in relation to water submersion. Briefly, our studies show that level of submersion, developmental stage, and species have profound impact on egg viability in relation to putative flooding events.

## Materials and Methods

### Tick egg species

Engorged female *D. albipictus* and *A. americanum* were obtained from the Knipling-Bushland US Livestock Insects Research Lab (KBUSLIRL; USDA-ARS, Kerrville, TX, USA). Tick colonies at this facility were initiated and are often supplemented with field-collected individuals and are maintained at 22 ± 1°C, 93% relative humidity (RH), and 14:10 hr light: dark condition (L:D). Both tick species are reared on beef cattle breeds (*Bos taurus*), and all animal procedures were approved by the Institutional Animal Care and Use Committee of the KBUSLIRL (#023-02, #023-03). Replete females were sent to our laboratory within 24 to 48 hours of host drop-off. Upon arrival, engorged females were held at 22 ± 1°C, 93% RH, and 14:10 hr L:D cycle until egg deposition (Winston and Bates 1960).

Some eggs were collected immediately following egg laying for early exposure (2-4 days after eggs had been laid), while others in the late exposure group were allowed to continue development (two weeks after eggs had been laid) before collection. Based on a previous assessment of the specific stages of embryogenesis, the eggs were in stages 4-5 of development (embryonic cells have moved to the periphery of the egg and have begun to proliferate) for early water submersion and stages 10-12 (presence of waste product in distinct sac-like structure and leg development can be observed) for late water submersion stress (Dipeolu, 1991; Santos et al., 2013).

### Water submersion experiments

Egg clusters, defined as the whole egg cluster deposited by a single female, were subjected to two treatments – half submersion and full submersion. As the egg clusters are hydrophobic, the half-submersion was accomplished by placing the entire egg cluster on the surface of the water (Bell mason jar, pint size). Full submersion was accomplished by placing the tick eggs in a mesh-covered container (Bell mason jar, pint size) that was held fully underwater within a large plastic container (Sterilte, 66 qt). The container remained 1-2 cm under the water. The ticks’ eggs were removed after 0 (control), 1, 2, 7, and 14 days and held at 93% RH for at least 12 weeks. Samples of the egg clusters and emerged larvae were frozen at -20 ± 1°C. The number of emerged larvae and eggs that did not hatch were counted three times independently.

### Statistical analyses

The proportion of emerged larvae (emerged larvae/total eggs treated) was used as the main indicator of difference in viability. Mean averages for all treatments are provided in Table S1. The proportion of eggs that hatched was calculated by dividing the number of hatched eggs by the total number of eggs for each treatment. We tested the effect of species, treatment, and developmental stage on the proportion of eggs that hatched and the timing of hatching using a linear regression model (lm function in the lme4 R package). For the initial analysis of broad patterns within the data, we started with models setting each response variable (i.e., proportion hatched) as a function of each predictor including egg development stage, species, and treatment. A residual analysis plot (residual values vs. expected values) was visually examined for all linear models to verify model assumptions of normality and homoscedasticity were met. Response variables were transformed (logit) as necessary to ensure that model assumptions were met. All additional analyses were performed using subsets of these models, including comparing within species, among treatments, and within development stages to allow for more targeted comparisons. Post-hoc comparisons were made using emmeans (default - Tukey method) where p < 0.05 were considered statistically significant. Output from models and post-hoc results for these linear models are shown in Supplemental Materials S1 and Supplemental Materials S2. All analyses were done in R version 3.6.3 (statistical packages - linear model - lme4, with data, with posthoc - emmeans) and data visualization was completed using tidyr, ggplot, and reshape2.

## Results

### General effects based on egg stage, submersion level, and tick species

Water immersion had a generally negative effect on *D. albipictus* and *A. americanum*, where both short period and prolonged exposures lead to a reduction of egg viability, (Fig. 1). When the egg stage was assessed, submersion during early egg development resulted in a more substantial reduction in egg viability (Fig. 1; F_1,142_=7.36, *P*=0.007). Submersion level also profoundly affected egg viability based on the time, where full submersion resulted in lower viability compared to those that experienced half submersion (Fig. 2; F_1,126_=4.731, *P*=0.006). Lastly, *A. americanum* eggs were much more tolerant to submersion stress than *D. albipictus* (Fig. 3; F_1,126_=68.59, *P*<0.001).

**Figure 1.**
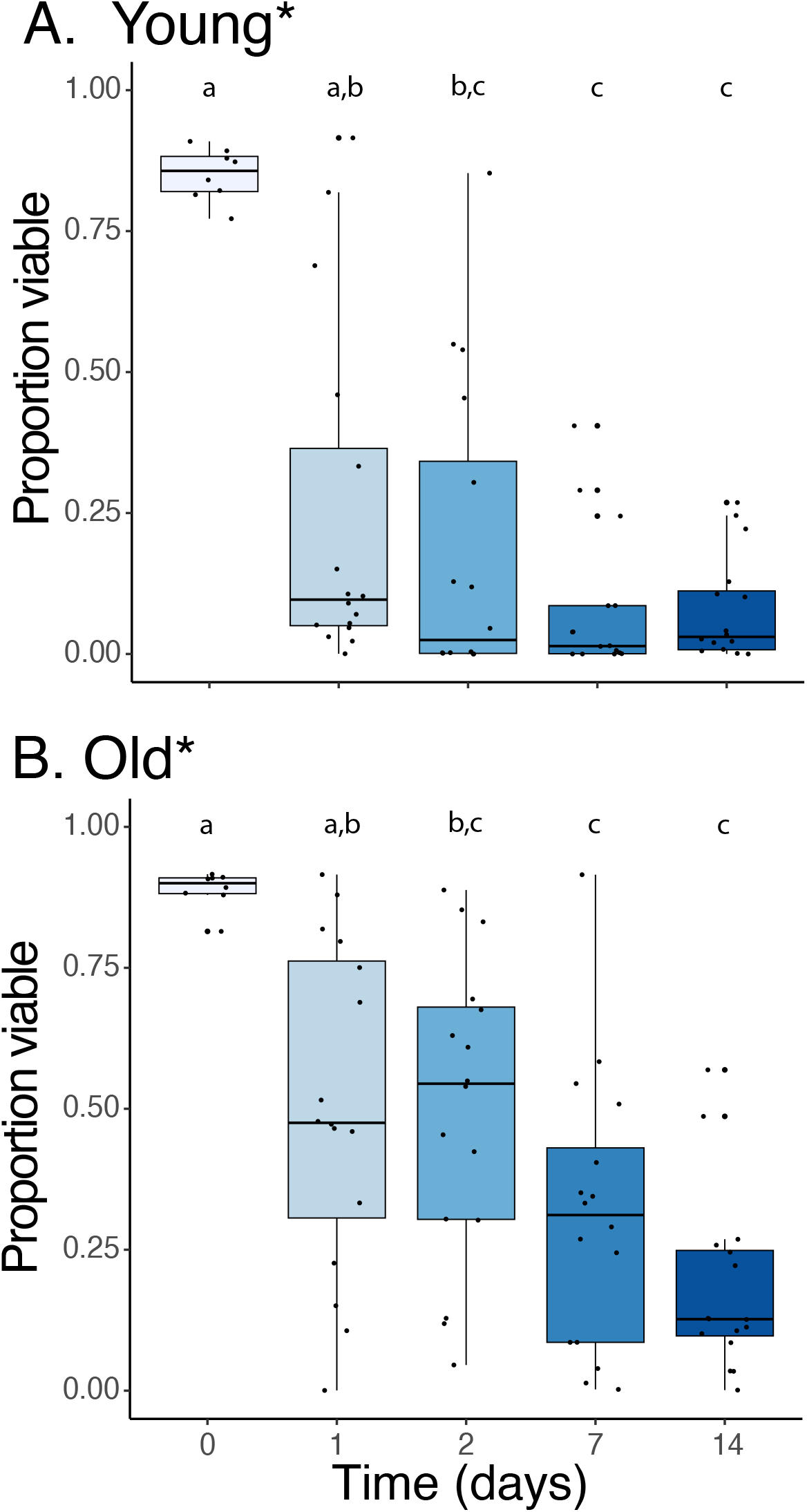
Impact of the developmental time on the viability of tick eggs in relation to water submersion. (A) Young tick eggs are characterized by the movement of the embryonic cells to the periphery of the eggs and have begun to proliferate and (B) old tick eggs are shown by the presence of waste product in a distinct sac-like structure and observable leg development (Dipeolu, 1991; Santos et al., 2013). * indicates a significant difference between survival between young and old tick eggs (F_1,142_=7.36, *P*=0.007). Letters indicate significant differences between groups (P < 0.05). Box plot whiskers show upper and lower bounds with the bottom of the boxes representing the bottom 25th percentile and the top 75th percentile. This data is pooled from two species: winter ticks, *Dermacentor albipictus*, and lone star ticks, *Amblyomma americanum*.

**Figure 2.**
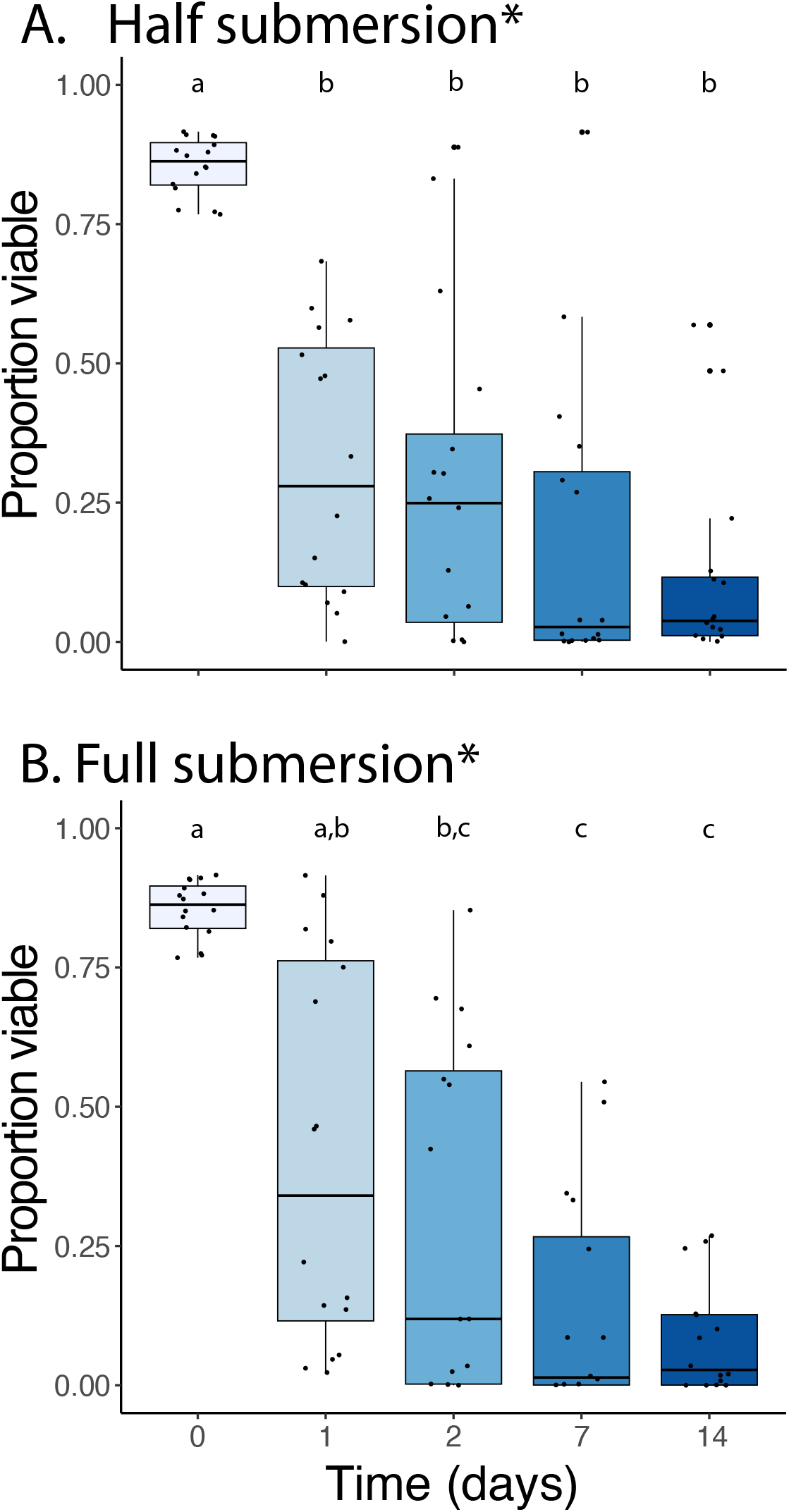
Effect of water submersion level on egg viability. (A) Half submersion (egg clutches floating on water) and (B) full submersion (egg clutches held under water). * indicates a significant difference between survival between full and half submersion (F_1,126_=4.731, *P*=0.006). Letters indicate significant differences between groups (P < 0.05). Box plot whiskers show upper and lower bounds with the bottom of the boxes representing the bottom 25th percentile and the top 75th percentile. This data is pooled from two species: winter ticks, *Dermacentor albipictus*, and lone star ticks, *Amblyomma americanum*.

**Figure 3.**
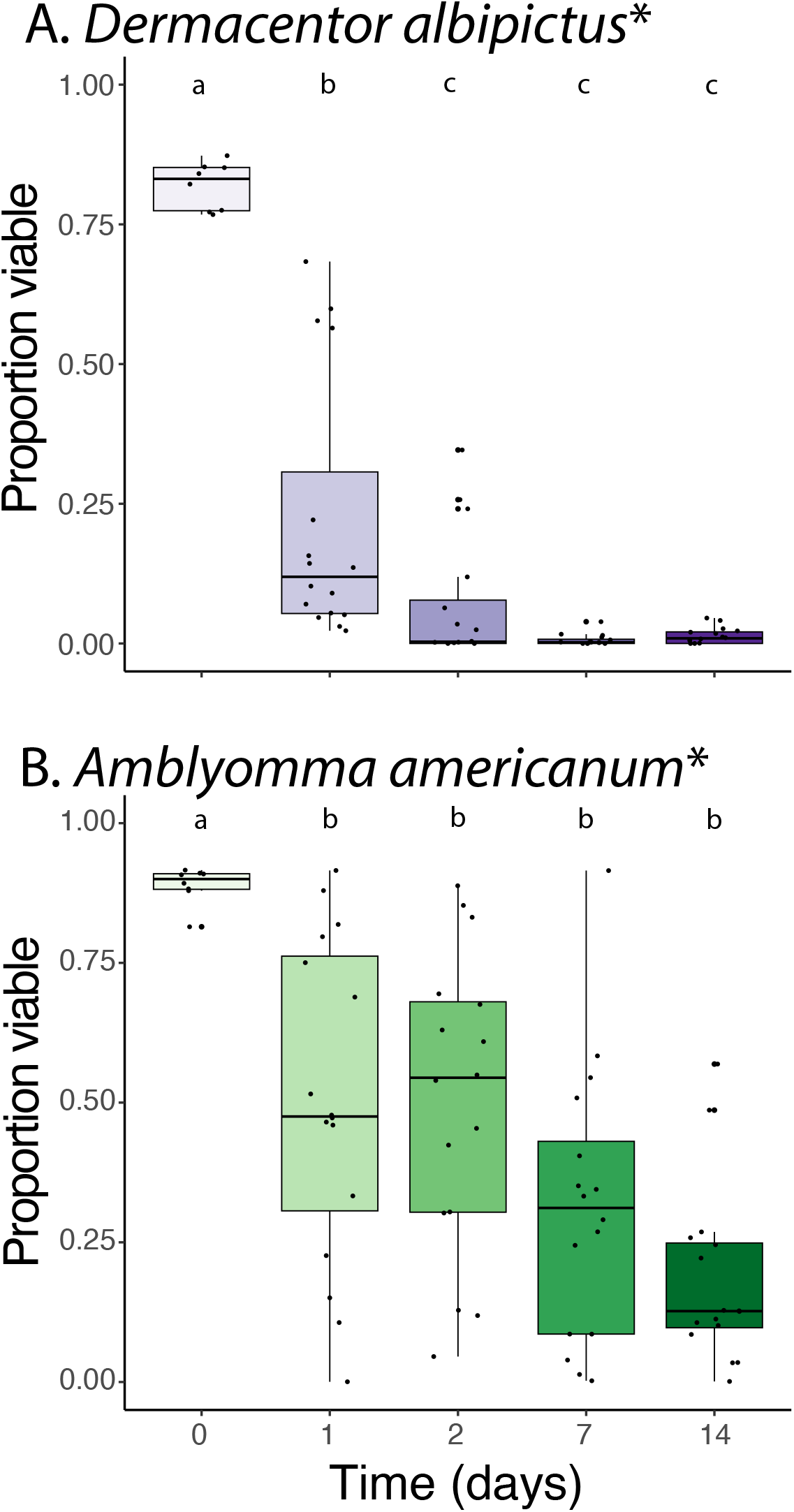
Species-specific differences in egg viability between winter ticks, *Dermacentor albipictus*, and lone star ticks, *Amblyomma americanum*. (A) *D. albipictus* and (B) *A. americanum*. * indicates a significant difference between survival between species (F_1,126_=68.59, *P*<0.001). Letters indicate significant differences between groups (P < 0.05). Box plot whiskers show upper and lower bounds with the bottom of the boxes representing the bottom 25th percentile and the top 75th percentile.

### Species-specific impact in relation to egg stage and submersion level

When *D. albipictus* was examined individually, there was a significant effect on the egg developmental stage, with eggs early in development having reduced viability compared to controls (Fig. 4A, 4B). The *D. albipictus* early-stage eggs showed nearly 80% reductions in viability after one and two days of submersion (half or full submersion (Fig. 4). When *D. albipictus* eggs were allowed to develop before water submersion, eggs remained viable, albeit 25-50% less than control groups with no water exposure. Prolonged periods of water submersion (7 and 14 days) yielded significant declines in viability for both young and old eggs.

**Figure 4.**
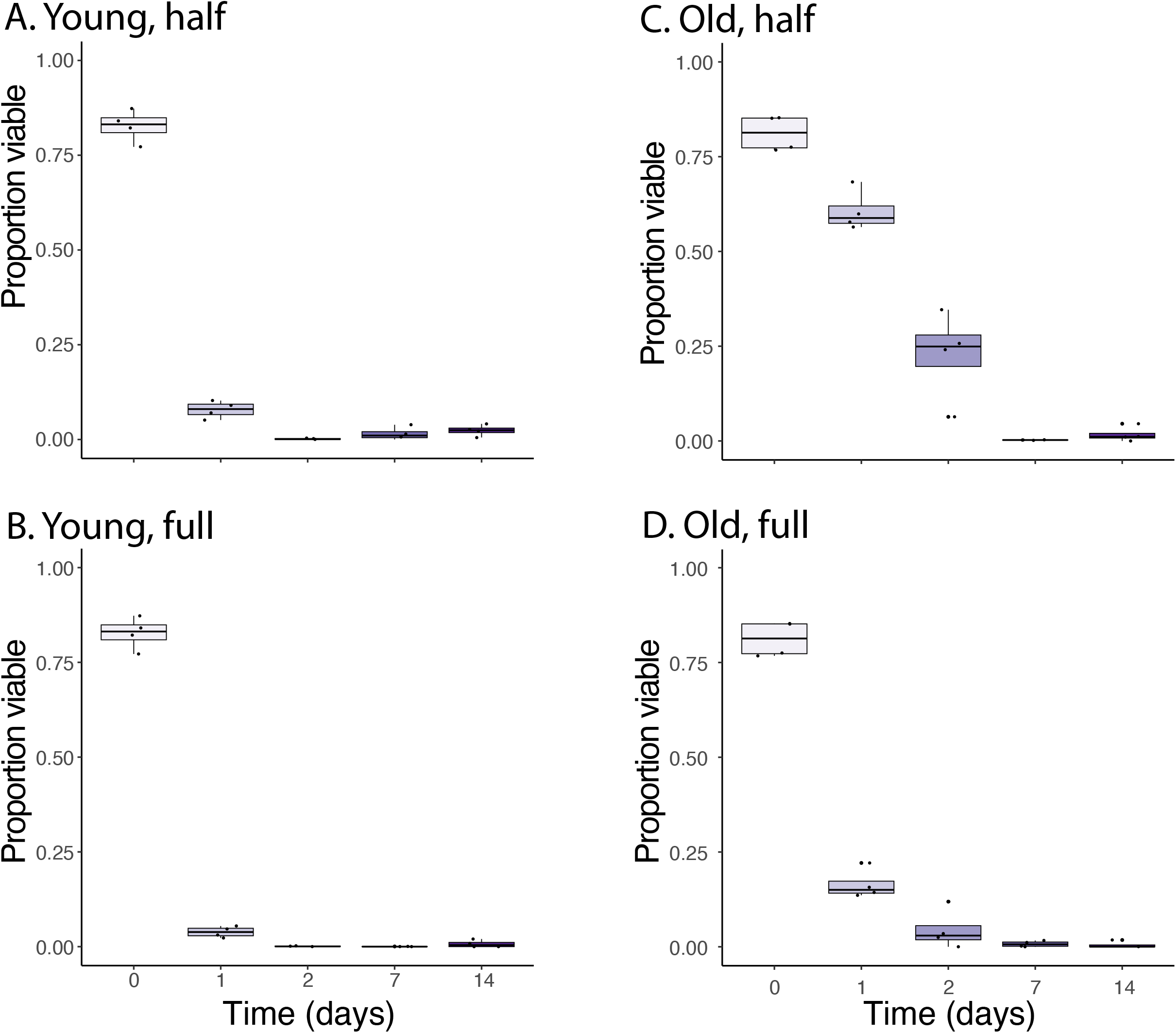
Egg viability of winter ticks, *Dermacentor albipictus*, in relation to egg developmental time and levels of water submersion. (A,B) Young tick eggs (Stages 4-5) and (C,D) old tick eggs (Stages 10-12). (A,C) Half submersion (egg clutches floating on water) and (B,D) full submersion (egg clutches held under water). Letters indicate significant differences between groups (P < 0.05). Box plot whiskers show upper and lower bounds with the bottom of the boxes representing the bottom 25th percentile and the top 75th percentile.

Compared to *D. albipictus, A. americanum* eggs were much more tolerant to water submersion (Fig. 5). Of interest, full versus half submersion only had a minor effect on *A. americanum* when exposed shorter periods (1 and 2 days); however, extended water exposure (7 and14 days) yielded reduced viability under full submersion compared to half submersion. As with *D. albipictus*, older eggs had much higher viability when submerged compared to less developed eggs (Fig. 5).

**Figure 5.**
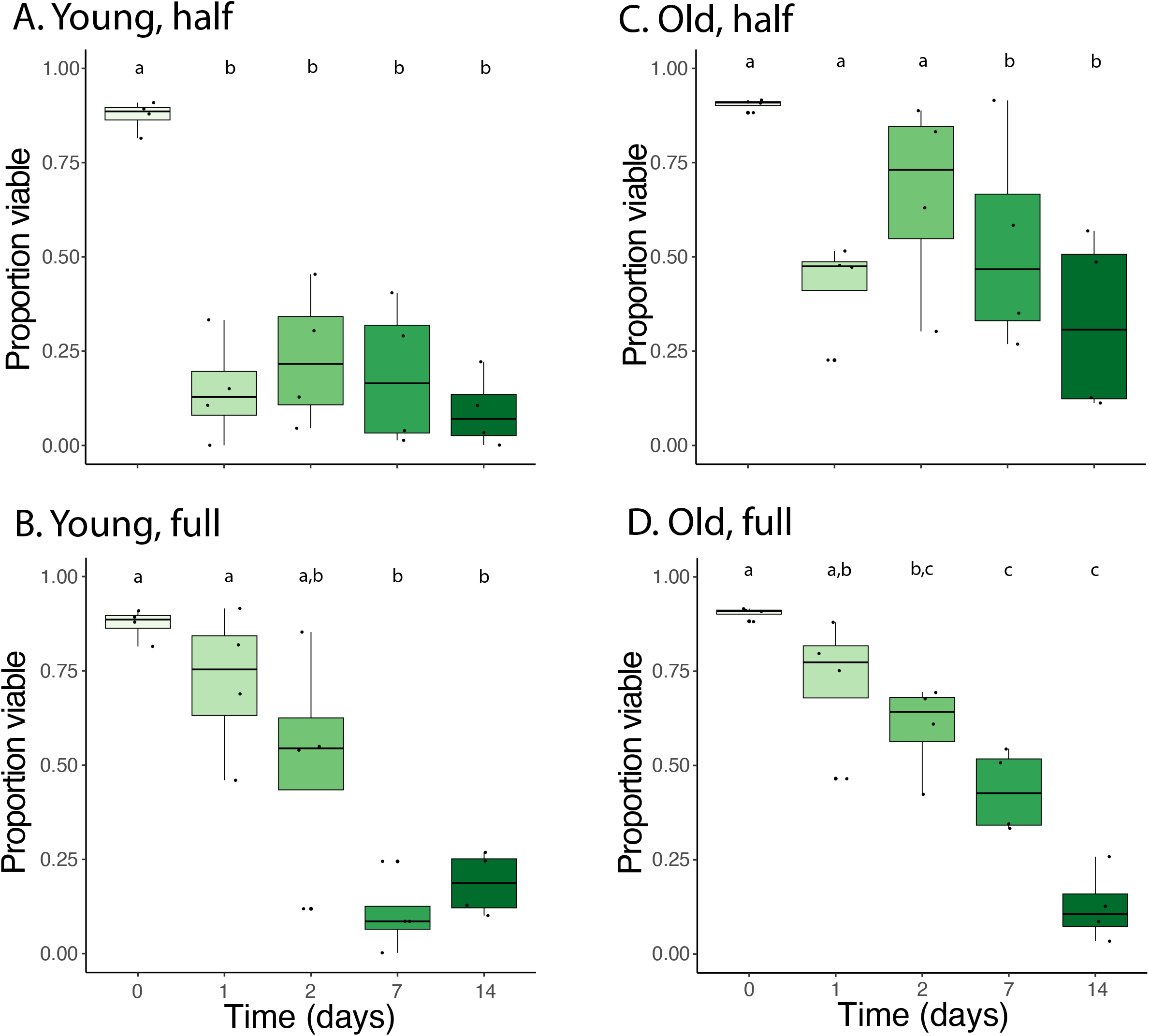
Egg viability of lone star ticks, *Amblyomma americanum*, in relation to egg developmental time and levels of water submersion. (A,B) Young tick eggs (Stages 4-5) and (C,D) old tick eggs (Stages 10-12). (A,C) Half submersion (egg clutches floating on water) and (B,D) full submersion (egg clutches held under water). Letters indicate significant differences between groups (P < 0.05). Box plot whiskers show upper and lower bounds with the bottom of the boxes representing the bottom 25th percentile and the top 75th percentile.

## Discussion

Here, we build upon a growing number of observations that flooding events will impact tick survival (Honzáková, 1971; Koch, 1986; Fielden et al., 2011; Giannelli et al., 2012; Bidder et al., 2019). Most of these studies focused on the mobile stages of ticks (larvae, nymphs, and adults). A few have assessed the viability of eggs, including *A. americanum* (Koch, 1986). Importantly, these studies examined eggs only under full submersion or focused on only a single developmental time point for eggs (Koch, 1986; Dipeolu, 1991; Santos et al., 2013). The results we obtained in these studies show the degree of submersion and the developmental stage of the eggs have profound effects on egg viability, and these differ between species.

As previously seen with thermal tolerance (Ajayi et al., 2024), the developmental stage of the egg had a major effect on survival after water submersion. Submersion in water was detrimental even after a single day. This impact was observed after two weeks of submersion in water for both *D. albipictus* and *A. americanum*. Along with the age of the eggs, the level of submersion had a significant impact on egg viability, where a partial submersion was less detrimental. Lastly, *D. albipictus* was less tolerant of water submersion compared to *A. americanum*. Altogether, these studies suggest that tick eggs are likely to remain viable after extended periods of water submersion, especially when the egg clutch is older and only partially submerged Furthermore, specific species are more tolerant to water submersion which likely translates to improved viability during flood exposure Our studies highlight that tick egg viability during periods of flooding should be considered when assessing the survival of tick species in specific areas.

## Acknowledgments

The authors thank Wayne Ryan (USDA-ARS) for assistance with tick rearing. Funding was provided by the University of Cincinnati as a Faculty Development Research Grant and, in part for dual usage of equipment, through the National Science Foundation DEB-1654417 and National Institute of Allergy and Infectious Diseases of the National Institutes of Health under Award Number R01AI148551, R21AI176098, and R21AI166633 to J.B.B.

